# Lymphatic ERG Governs Junctional Plasticity and Fluid Clearance via an EDNRB Signaling Axis in Pulmonary Fibrosis

**DOI:** 10.64898/2026.09.17.752506

**Authors:** Arun Narota, Tapiwa Muvavarirwa, Xintao Qiu, Adri Chakraborty, Ahmed A. Raslan, Giovanni Ligresti, Maria Trojanowska

## Abstract

Pulmonary lymphatic vessels are essential for interstitial fluid balance and macromolecular clearance, yet the endothelial mechanisms regulating lymphatic vessel function during chronic fibroproliferative lung injury remain undefined. Here, we demonstrate that the lineage-defining transcription factor ERG in lymphatic endothelial cells (LECs) governs junctional plasticity and fluid drainage in pulmonary fibrosis. In human idiopathic pulmonary fibrosis (IPF) lungs, lymphatic ERG expression is markedly downregulated. Lineage-specific inducible deletion of *Erg* in murine LECs (Erg-CKO) unexpectedly conferred robust protection against bleomycin-induced pulmonary fibrosis, significantly dampening acute inflammation, reducing edema, and preserving pulmonary compliance. Mechanistically, loss of ERG enhanced lymphatic drainage capacity in vivo. Transcriptomic profiling of isolated LECs revealed that ERG deficiency activates actomyosin contractile pathways and selectively upregulates Endothelin Receptor Type B (*Ednrb*), driving junctional remodeling from continuous zippers into discontinuous button-like configurations that facilitate interstitial fluid entry. Pharmacological blockade of EDNRB with BQ-788 abolished junctional reorganization, eliminated the enhanced lymphatic drainage, and completely reversed the anti-fibrotic protection in Erg-CKO mice. Together, our findings identify an endothelial ERG-EDNRB regulatory axis that controls lymphatic junctional architecture and demonstrate that promoting lymphatic clearance via endothelial junctional remodeling represents a viable therapeutic strategy for fibrotic vascular remodeling.

## INTRODUCTION

Idiopathic Pulmonary Fibrosis (IPF) is characterized by irreversible architectural destruction of lung parenchyma, driving terminal respiratory failure (1, 2). A central feature of early acute lung injury that sets the stage for progressive fibrosis is the destabilization of the alveolar-capillary barrier (3). This breach causes an accumulation of protein-rich edema, inflammatory cytokines, and cellular debris within the interstitial space. Under normal physiological conditions, the pulmonary lymphatic network serves as the primary drainage route, maintaining fluid homeostasis and removing tissue debris to restore tissue architecture (4).

Unlike peripheral or mesenteric collecting lymphatics, which are enveloped by continuous layers of smooth muscle cells (SMCs) that drive active pumping, collecting lymphatics within the delicate lung parenchyma operate in a highly compliant mechanical environment and lack SMC coverage. Consequently, pulmonary lymphatic endothelial cells (LECs) do not rely on extrinsic muscular pumping. Instead, they depend on intrinsic actin-myosin contractility, rapid junctional adaptation, and local mechanical deformation driven by respiratory movement to drain fluid into the regional mediastinal lymph nodes (mLNs) (5, 6).

The transcription factor ERG (ETS-related gene) is well known as an essential master regulator of endothelial cell homeostasis. In the vascular endothelium, ERG maintains junctional integrity, primarily through the transcription of VE-cadherin, and prevents vascular hyperpermeability, inflammation, and thrombosis (7–9). Targeted loss of ERG in vascular endothelial cells causes junctional breakdown, excessive microvascular leakage, and tissue inflammation (10). While prior studies using pan-endothelial knockouts have established a core role for ERG in vascular biology, these observations primarily reflect blood endothelial cells (BECs) due to their sheer abundance relative to LECs. Consequently, whether ERG plays a specialized role in mature LECs, particularly within the distinct structural microenvironment of the lung, remains unknown.

In this study, we investigated the cell-type-specific function of ERG in adult pulmonary lymphatics during fibroproliferative lung injury. To address this, we used conditional genetic deletion of ERG specifically within adult mouse LECs (*Prox1*Cre*ERT2;Erg^fl/fl^*; Erg-CKO) and demonstrated that targeting ERG confers protection against bleomycin-induced pulmonary fibrosis. We demonstrate that ERG loss triggers an upregulation of the endothelin B receptor (EDNRB), which significantly enhances lymphatic drainage and accelerates the clearance of pro-inflammatory cytokines, excess fluid (edema), and damaged cells from the lung. These results identify the ERG-EDNRB axis as a target that can be manipulated to enhance lymphatic function and prevent fibrotic progression.

## RESULTS

### Loss of lymphatic ERG expression is a hallmark of pulmonary fibrosis

To investigate the translational relevance of ERG in patients with IPF, we first analyzed its expression in human single-cell RNA sequencing (scRNA-seq) datasets from patients with IPF (**Fig. 1a**). Across multiple independent patient cohorts (11–13), *ERG* transcript levels in LECs were consistently and markedly lower in IPF lungs than in healthy controls (HCs; **Fig. 1a**).

**Figure 1.**
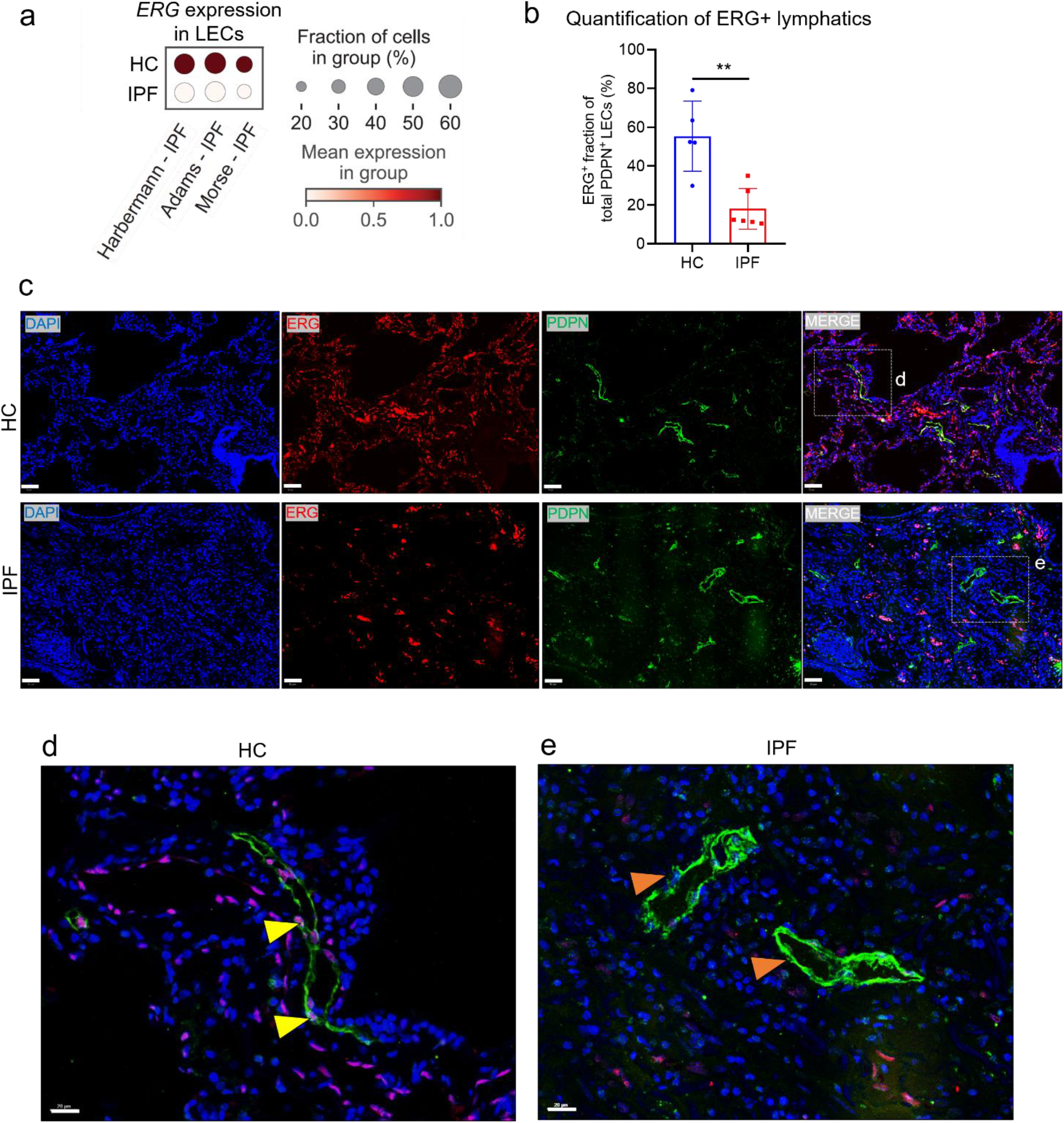
Lymphatic ERG expression is downregulated in human IPF. **a)** Dot plot showing single-cell RNA sequencing meta-analysis of *ERG* expression in pulmonary LECs across three independent human datasets (Habermann, Adams, and Morse) comparing HC and IPF lungs. Dot size indicates the fraction of cells expressing *ERG* (%), and color intensity reflects the mean expression level within the group. **b)** Quantification analysis of ERG+ LECs expressed as ERG+ fraction of total PDPN+ LECs (%) in lung sections from HCs (n=4) and patients with IPF (n=6). **c)** Representative IF images of lung tissues form HC (top panel) and patients with IPF (bottom panel) stained for DAPI (nuclei, blue), ERG (red), and PDPN (lymphatic endothelium, green), along with merged channels. Scale bars = 50µm. **d, e)** High-magnification views of the boxed regions in preceding panel highlighting d) nuclear ERG expression in PDPN+ lymphatic endothelial cells of HC lungs (yellow arrowheads) versus e) loss of ERG nuclear signal in PDPN+ lymphatic vessels of IPF lungs (orange arrowheads). Scale bars = 20 µm.

To validate these transcriptomic findings at the protein level, we performed immunofluorescence (IF) staining on human lung tissue sections from HCs and IPF patients, using Podoplanin (PDPN) as a specific marker for lymphatic vessels and ERG to mark endothelial nuclei (**Fig. 1c**). Quantitative analysis of these sections revealed a significant reduction in the fraction of ERG-positive lymphatics in IPF tissues compared with that in healthy tissues, from ∼55% in HCs to below 20% in IPF tissues (**p<0.01; **Fig. 1b**). High-magnification imaging of healthy lungs showed bright nuclear localization of ERG within the endothelial wall of PDPN+ lymphatic vessels (**Fig. 1d**, yellow arrowheads). Conversely, lymphatic vessels in the fibrotic regions of IPF lungs displayed a striking loss of nuclear ERG signal despite maintaining structural PDPN positivity (**Fig. 1e**, orange arrowheads). Together, these clinical data demonstrate that lymphatic ERG expression is downregulated during human pulmonary fibrosis, suggesting that its absence may be an active contributor in directing lymphatic dysfunction in the pathogenesis of IPF.

### Loss of lymphatic ERG attenuates bleomycin-induced pulmonary fibrosis in mice

Having observed a downregulation of lymphatic ERG expression in human IPF lungs, we next examined the functional consequences of lymphatic ERG loss *in vivo* using an inducible, conditional knockout mouse model (*Prox1*Cre/*Erg*^fl/fl^, hereafter Erg-CKO). Following tamoxifen induction and a three-week washout period, mice were subjected to intratracheal (*i.t.*) bleomycin administration to induce pulmonary fibrosis (**Fig. 2a**). Lungs were harvested at 14 days post-injury (14dpi), which corresponds to the peak of the fibrotic response, to assess tissue remodeling and extracellular matrix (ECM) deposition (**Fig. 2a**).

**Figure 2.**
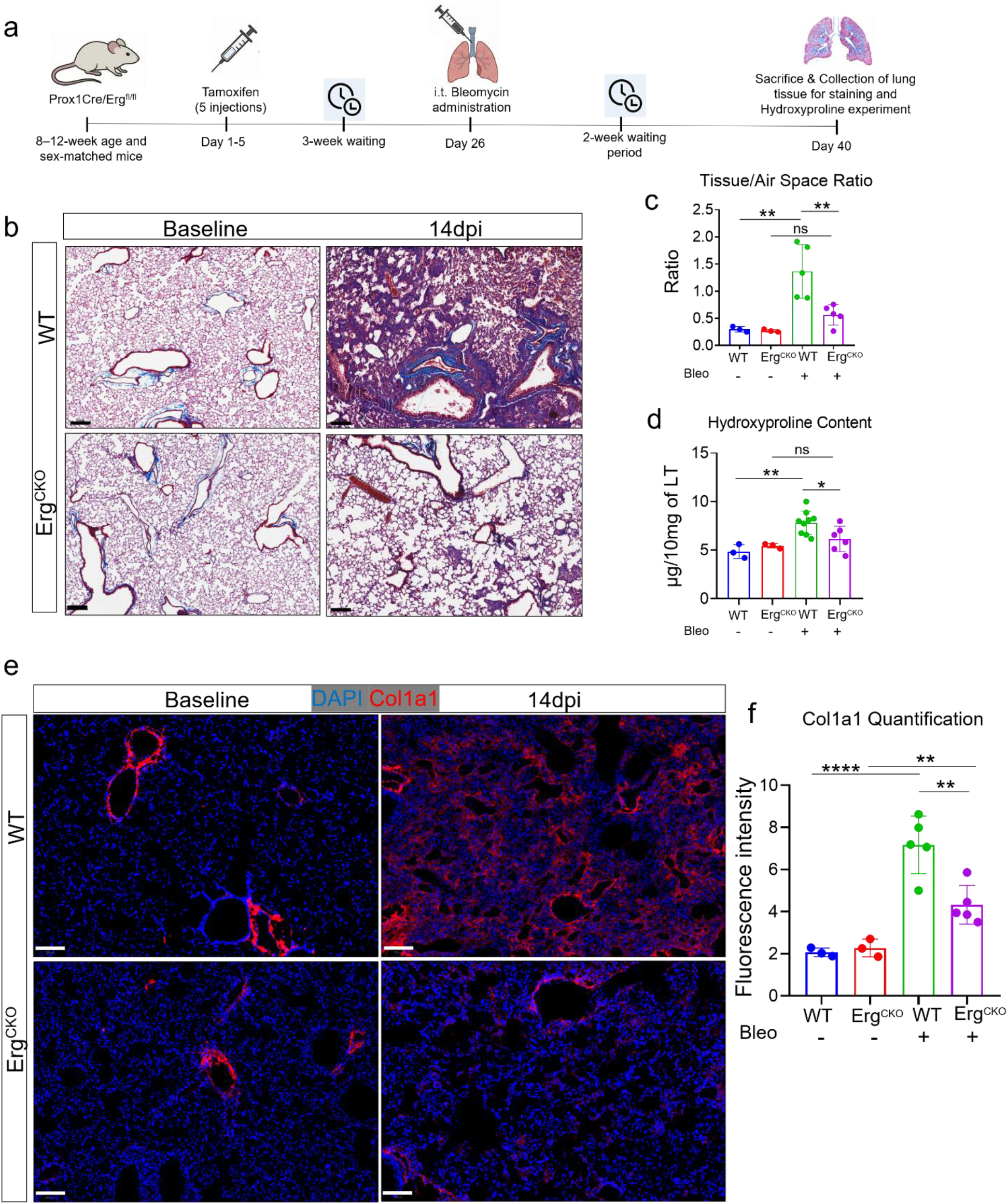
Lymphatic ERG deletion attenuates experimental lung fibrosis in mice. **a)** Schematic experimental timeline for tamoxifen-induced lymphatic endothelial cell (LEC)-specific deletion of Erg (Prox1-CreERT2;Erg^fl/fl^, termed Erg-CKO) and subsequent *i.t.* bleomycin administration (harvested at 14dpi). **b)** Representative images of Masson’s trichrome staining of lung sections from WT and Erg-CKO mice at baseline and 14dpi. Scale bars = 200 µm. **c)** Morphometric quantification of parenchymal fibrosis expressed as the tissue-to-airspace ratio in lung sections from PBS-and bleomycin-administered WT and Erg-CKO mice (n=3–5). **d)** Total lung collagen deposition quantified by hydroxyproline assay (µg/10 mg of lung tissue) across experimental groups (n=3–9). **e)** Representative IF staining for Col1a1 (red) and nuclei (DAPI, blue) in lung sections from WT and Erg-CKO mice under baseline conditions and at 14dpi. Scale bars = 100 µm. **f)** Quantitative fluorescence intensity analysis of Col1a1 immunoreactivity in lung sections across experimental groups (n=3–5).

Under baseline conditions, histological examination via Masson’s Trichrome staining revealed no baseline architectural or collagen deposition differences between wild-type (WT) and Erg-CKO mice (**Fig. 2b**). However, at 14dpi, bleomycin-treated WT mice exhibited severe parenchymal remodeling and dense collagen deposition (blue staining) across the lung parenchyma (**Fig. 2b**). Surprisingly, this pathological remodeling was substantially attenuated in Erg-CKO mice, which maintained relatively preserved alveolar structures and showed noticeably less collagen and ECM accumulation (**Fig. 2b**). Morphometric analysis of the tissue-to-airspace ratio confirmed that the architectural damage observed in injured WT lungs was significantly reduced in the Erg-CKO group (**p<0.01; **Fig. 2c**). To biochemically quantify total collagen content, we performed a hydroxyproline assay. Consistent with our histological observations, bleomycin injury caused a robust increase in hydroxyproline levels in WT lungs (**p<0.01), which was significantly blunted in those from Erg-CKO mice (*p<0.05; **Fig. 2d**).

To further evaluate ECM production at the molecular level, we performed IF staining for Collagen Type I Alpha 1 (Collagen-I) on whole lung sections (**Fig. 2e**). While baseline Collagen-I expression was low and restricted primarily to the perivascular and peribronchial niches across both genotypes, bleomycin injury triggered upregulation of interstitial Collagen-I throughout the parenchymal field in WT mice (**Fig. 2e**). In contrast, Erg-CKO mice displayed an attenuated increase Collagen-I expression post-injury (**Fig. 2e**). Quantitative analysis of the fluorescence intensity confirmed that interstitial Collagen-I deposition was significantly lower in the Erg-CKO group than injured WT controls (**p<0.01; **Fig. 2f**). Taken together, these data demonstrate that the selective loss of lymphatic ERG provides protection against experimental pulmonary fibrosis, reducing both structural parenchymal destruction and collagen deposition.

### Loss of lymphatic ERG expression blunts the acute inflammatory response and attenuates pulmonary edema after bleomycin injury

Following our observation that Erg-CKO mice are protected from bleomycin-induced fibrosis, we sought to determine if this protection was rooted in an altered early inflammatory and edematous response. We utilized a standard experimental timeline where WT and Erg-CKO mice were administered *i.t.* bleomycin and analyzed at various time points (3, 7, and 14dpi; **Fig. 3a**). We quantified inflammatory cell infiltration, IL-6 levels, and total protein content by analyzing Bronchoalveolar Lavage Fluid (BALF). Bleomycin challenge in WT mice led to a time-dependent influx of immune cells into the alveolar space, peaking at 7dpi, as evidenced by a significant increase in total cell number and IL-6 protein levels (**Fig. 3b-c)**. Moreover, Erg-CKO mice demonstrated a significant reduction in total inflammatory cell influx into the BALF at 7dpi compared to WT mice (****p<0.0001), though cell numbers remained slightly elevated above baseline levels (**Fig. 3b**). Further, bleomycin challenge in WT mice triggered a robust, time-dependent surge in BALF IL-6 levels that peaked at 7dpi before declining toward baseline by 14dpi (**Fig. 3c**). Reflecting the broader reduction in inflammation, Erg-CKO mice exhibited a marked suppression of this IL-6 burst, showing significantly lower levels at both 7dpi and 14dpi compared with WT controls (****p<0.0001; **Fig. 3c**). We next assessed pulmonary edematous changes, which are a hallmark of acute lung injury. Consistent with inflammatory cell influx, the Lung wet/dry weight ratio, a standard measure of pulmonary fluid accumulation, was significantly increased in WT mice at 7dpi compared to baseline levels (***p<0.001), indicating substantial edema (**Fig. 3d**). In contrast, Erg-CKO mice exhibited a marked reduction in this edematous response, with wet/dry ratios at 7dpi remaining significantly lower than in WT controls (***p<0.001) and closer to baseline levels (**Fig. 3d**). This attenuation of pulmonary fluid accumulation at 7dpi suggests that lymphatic ERG deletion enhances parenchymal fluid clearance during acute phase of injury. Furthermore, total BALF protein content, an indicator of alveolar-capillary barrier breakdown, was also significantly reduced in Erg-CKO mice at 7dpi compared to WT mice (***p<0.001; **Fig. 3e**). Together, these results demonstrate that deleting Erg specifically in the adult lymphatic endothelium leads to a blunted inflammatory and edematous response following bleomycin injury. This early attenuation of acute inflammation and edema may contribute to the reduced severity of subsequent chronic pulmonary fibrosis observed in this model.

**Figure 3.**
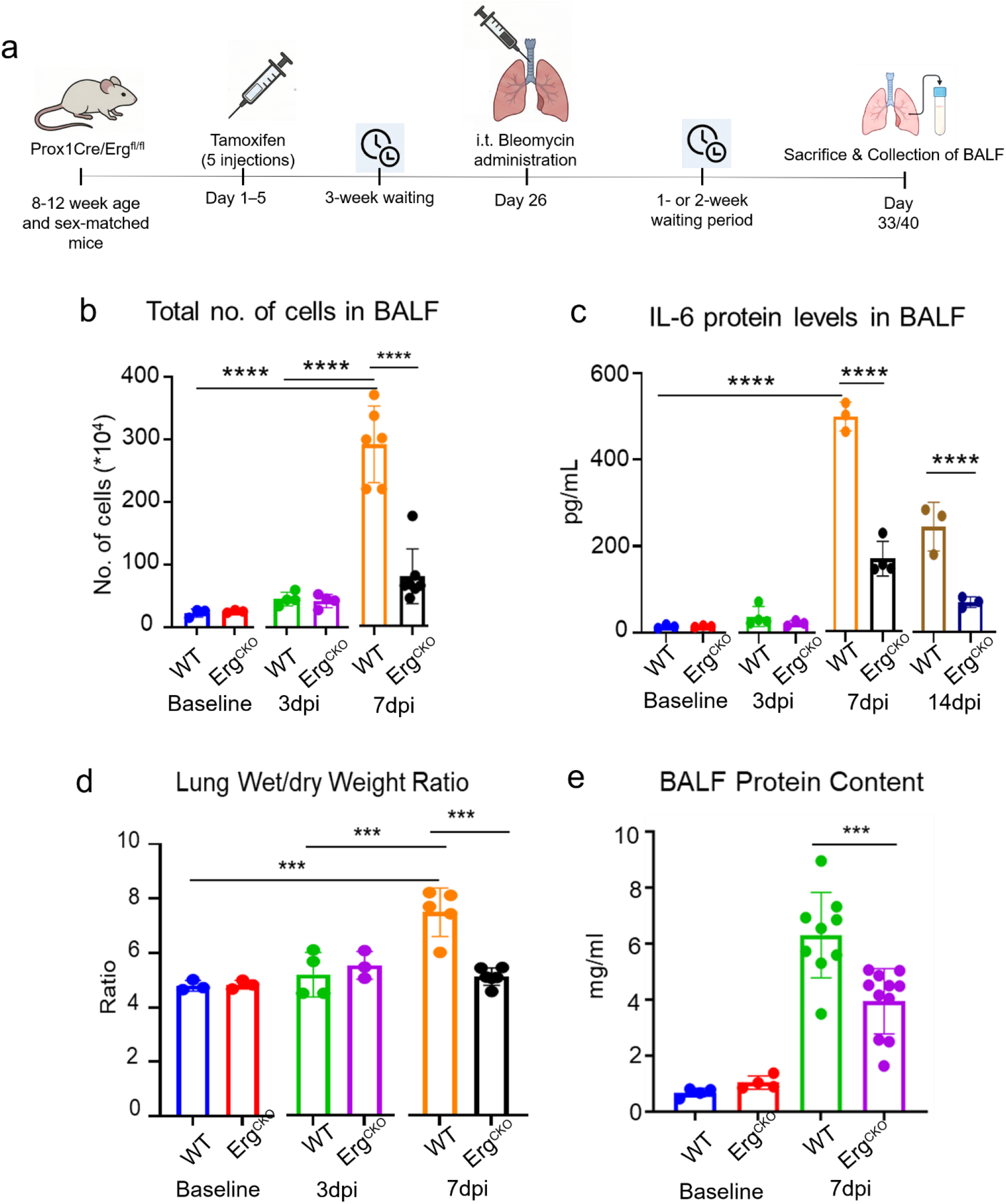
Lymphatic ERG deletion dampens inflammation in bleomycin-induced lung injury. **a)** Schematic experimental timeline illustrating the induction of Erg-CKO, followed by *i.t.* bleomycin instillation and BALF collection during the acute inflammatory phase (up to 7dpi or 14dpi). **b)** Total inflammatory cell count (10^4^ cells) quantified in BALF from WT and Erg-CKO mice at baseline, 3dpi, and 7dpi (n=3–6). **c)** IL-6 protein concentrations (pg/mL) measured by ELISA in BALF of WT and Erg-CKO mice across baseline, 3dpi, 7dpi, and 14dpi timepoints (n=3–4). **d)** Assessment of interstitial edema determined by lung wet-to-dry weight ratio in WT and Erg-CKO mice under baseline conditions, 3dpi, and 7dpi (n=3–5). **e)** Total protein concentration (mg/mL) in BALF from WT and Erg-CKO mice under baseline conditions and 7dpi (n=4–11).

### Loss of lymphatic ERG enhances pulmonary drainage independent of lymphangiogenesis

To determine if the attenuated inflammatory and fibrotic responses in Erg-CKO mice were linked to improved lymphatic function, we performed an *in vivo* drainage assay using high-molecular-weight FITC-dextran (150 kDa). Following *i.t.* administration, we quantified the transport of the tracer from the lungs to nearest mLNs, which serve as the primary drainage site for the lung parenchyma (**Fig. 4a**). Interestingly, Erg-CKO mice exhibited significantly higher lymphatic drainage capacity compared to WT mice under baseline conditions (***p<0.001; **Fig. 4b, c**). This enhanced drainage was maintained at 7dpi, with Erg-CKO mLNs showing nearly double the fluorescence intensity of their WT counterparts (**Fig. 4c**). These functional data suggest that the loss of Erg primes the lymphatic system for more efficient fluid and macromolecule clearance, potentially explaining the reduced pulmonary edema observed in Figure 3. We next investigated whether this functional shift was accompanied by structural changes in the lymphatic network. We performed IF staining for GFP across whole lung sections utilizing WT (*Erg*^+/+^*Prox1*-CreERT2^+^tdTomato^+^) and Erg-CKO GFP (*Erg*^fl/fl^*Prox1*-CreERT2^+^tdTomato^+^) reporter mice to assess lymphatic vessel density at baseline and 7dpi time points (**Fig. 4d**). Quantitative analysis revealed that while vessel density (vessel area/total lung area) was comparable between genotypes at both the time points (baseline and 7dpi), there was an increase in vessel density as the injury progressed after bleomycin administration (**Fig. 4e**). Thus, the enhanced drainage capacity observed in Erg-CKO mice was not a result of increased vessel density or de novo lymphangiogenesis. Instead, the loss of ERG appears to promote a more efficient drainage mechanism for the clearance under baseline and injury-induced stress.

**Figure 4.**
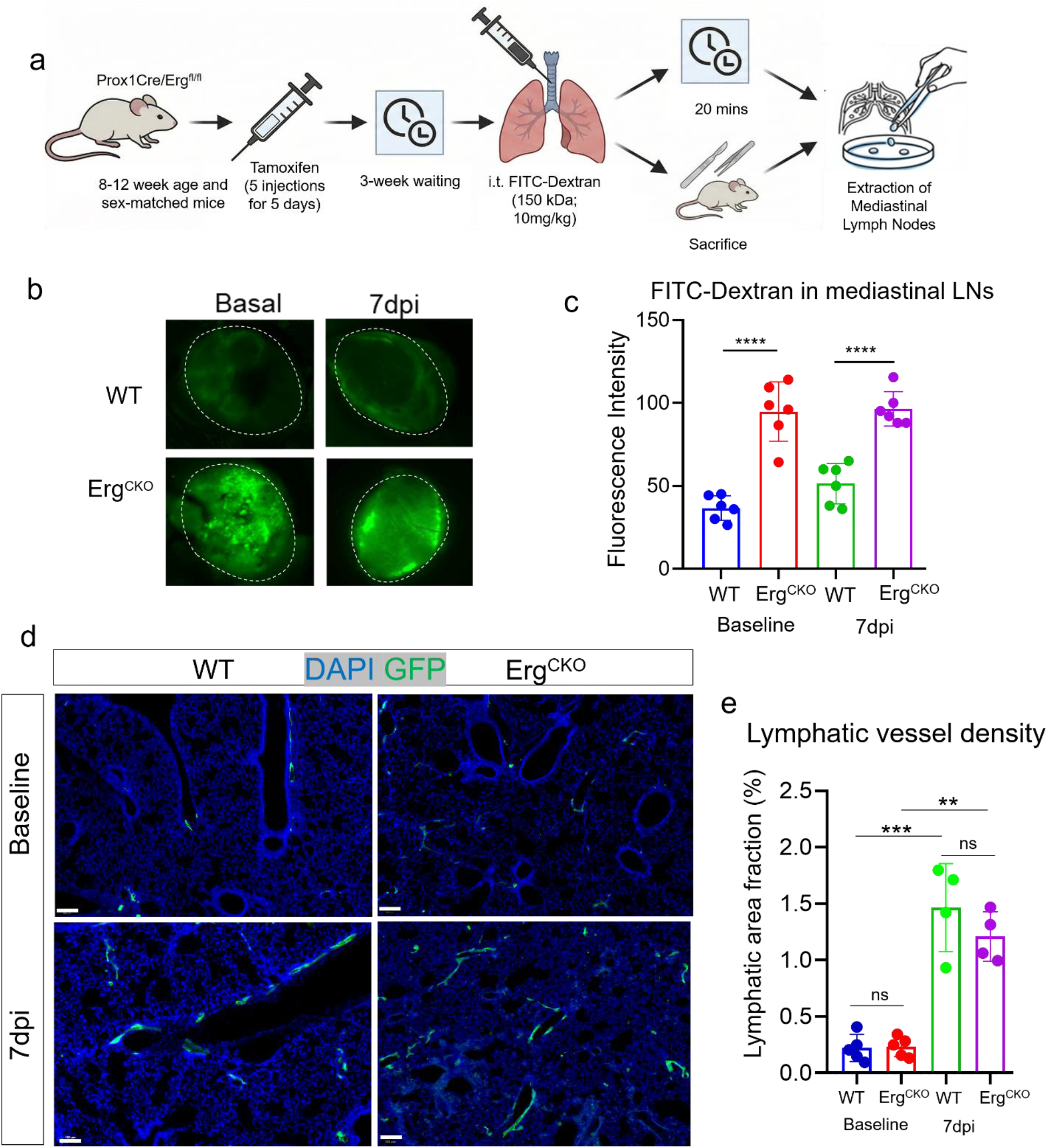
Lymphatic ERG deletion enhances drainage efficiency, and this mechanism is independent of lymphangiogenesis. **a)** Experimental workflow for assessing pulmonary lymphatic drainage kinetics in WT and Erg-CKO mice. **b)** Representative fluorescence images showing FITC-Dextran accumulation within regional draining mLNs (dashed outlines) from WT and Erg-CKO mice under baseline conditions and at 7dpi. **c)** Quantification of mean fluorescence intensity of accumulated FITC-Dextran in draining mLNs across experimental groups (n=6). **d)** Representative IF confocal images of lung sections stained for endogenous lymphatic reporter expression (GFP, green) and nuclei (DAPI, blue) in WT and Erg-CKO mice at baseline and 7dpi. Scale bar = 100µm. **e)** Morphometric quantification of pulmonary lymphatic vessel density expressed as the lymphatic area fraction (%) across lung sections in WT and Erg-CKO mice at baseline conditions and 7dpi (n=4–5).

### Transcriptomic profiling reveals that lymphatic *Erg* loss upregulates *Ednrb* and *Hgf*

To elucidate the molecular mechanisms underlying the enhanced drainage and protective phenotype of Erg-CKO mice, we performed bulk RNA-sequencing on primary LECs isolated from adult mouse lungs. Using a tdTomato-GFP (*Erg*^fl/fl^*Prox1*-CreERT2^+^tdTomato^+^) reporter system, we enriched for Cd31+ endothelial cells via MACS, followed by FACS sorting for GFP+ LECs (**Fig. 5a, b**). Principal component analysis (PCA) of global transcriptomic profiles revealed distinct segregation between baseline WT and Erg-CKO LECs along both principal components (PC1: 53.4%, PC2: 46.6%), confirming clear transcriptional divergence driven by lymphatic ERG deficiency (**Supplementary Fig. 3b**). Heatmap analysis demonstrated a broad and consistent downregulation of canonical lymphatic endothelial identity genes in Erg-CKO LECs compared to WT controls (**Supplementary Fig. 3c**) as previously observed in dermal lymphatics (14). Concordant with our sequencing data, RT-qPCR validation confirmed significant reductions in *Lyve1* (*p<0.05) and *Flt4* (**p<0.01) expression in sorted GFP^+^ Erg-CKO LECs compared to WT controls, while *Prox1* transcript levels showed no significant change (**Supplementary Fig. 3d**). Along with expected loss of *Erg*, we observed reduced expression of core lymphatic markers including *Nrp2, Mmrn1, Pdpn, Lyve1, Flt4,* and *Sox18* (**Supplementary Fig. 3c**). Ingenuity Pathway Analysis (IPA) of differentially expressed genes revealed significant enrichment (Z-score>2.0) for pathways governing actomyosin contractility, Rho family GTPases, actin cytoskeleton signaling, and motility, alongside concurrent inhibition of the RHOGDI inhibitory pathway in baseline Erg-CKO LECs (**Fig. 5c**). In line with this pathway-level activation, heatmap analysis of specific contractility-associated genes identified via IPA demonstrated coordinated upregulation of key cytoskeletal and contractile genes in Erg-CKO LECs, including striated and smooth muscle-related actins (*Acta1*, *Actc1*), myosins (*Myl1*, *Myl4*, *Myl7*, *Myh1*, *Myh6*, *Myh8*), troponins (*Tnni3*, *Tnnc2*, *Tnnt3*), and tropomyosins (*Tpm1*, *Tpm2*), alongside selective modulators like *Rnd3* and *Pak3* (**Supplementary Fig. 3e**). Further, consistent with this activated, contractile phenotype, transcriptomic profiling highlighted a selective induction of genes associated with endothelial barrier permeability, junctional dynamics, and paracrine signaling (**Fig. 5d**). Key regulators of endothelial remodeling and shear stress response, including *Gata2, Gja4, Angpt2,* and *Klf2*, were concurrently upregulated, while the junctional protein *F11r* (JAM-A) was downregulated in Erg-CKO LECs (**Fig. 5d**). Crucially, the Endothelin receptor B (*Ednrb*) and Hepatocyte Growth Factor (*Hgf*) were among the most significantly upregulated genes in Erg-CKO LECs (**Fig. 5d**). Notably, we observed similar trends in the lymphatic permeability-associated genes in siERG-treated human pulmonary LECs, including the upregulation of *EDNRB*, *HGF*, *GATA2*, and *GJA4* and downregulation of *CDH5*, though *KLF2* showed differential regulation relative to mouse LECs (**Supplementary Fig. 4a**). To confirm our transcriptomic findings, we performed independent validation via RT-qPCR on mouse-isolated LECs (**Fig. 5e**). We confirmed a reduction in *Erg* mRNA (**p<0.01), accompanied by significant fold-change increases in both *Ednrb* (**p<0.01) and *Hgf* (*p<0.05) mRNA in Erg-CKO LECs compared to WT LECs (**Fig. 5e**). Because HGF is a secreted anti-fibrotic factor, we quantified circulating protein levels in peripheral blood. Serum ELISA demonstrated significantly higher circulating HGF protein concentrations in Erg-CKO mice relative to WT controls (*p<0.05; **Fig. 5f**). We were particularly intrigued by the upregulation of *Ednrb,* because of its known role in vascular tone regulation (15). To determine if endothelin/EDNRB pathway is relevant to human disease, we analyzed publicly available IPF datasets (11–13). Analysis of *EDNRB* expression in healthy human lungs revealed the highest level of expression in aerocytes, while lower levels were detected in lymphatics, pericytes and smooth muscle cells. In terms of *EDNRB* ligands, *EDN1* is the most abundant isoform in the lungs and is expressed in artery, capillary and vein cells (**Fig. 5g**). In patients with IPF, there was overall reduced level of *EDNRB* expression across all cell types, while expression of EDN1 was increased (**Fig 5g**). Specifically, LEC expression of the *EDNRB* gene in patients with IPF showed an ∼38% decrease compared to HCs (adjusted p<0.001; **Fig. 5g**). This clinical trend was recapitulated in our WT mouse model, where lung *Ednrb* transcript showed a declining trend at 7 and 14dpi (**Supplementary Fig. 4c**). This suggests that although ERG loss increases EDNRB expression, other factors within the inflammatory and fibrotic milieu may counteract this effect, resulting in reduced EDNRB levels in both human IPF lungs and mouse models of pulmonary fibrosis. Collectively, these data demonstrate that ERG deficiency downregulates classic lymphatic identity markers and drives the upregulation of *Ednrb,* establishing a transcriptional state associated with enhanced lymphatic function and tissue protection.

**Figure 5.**
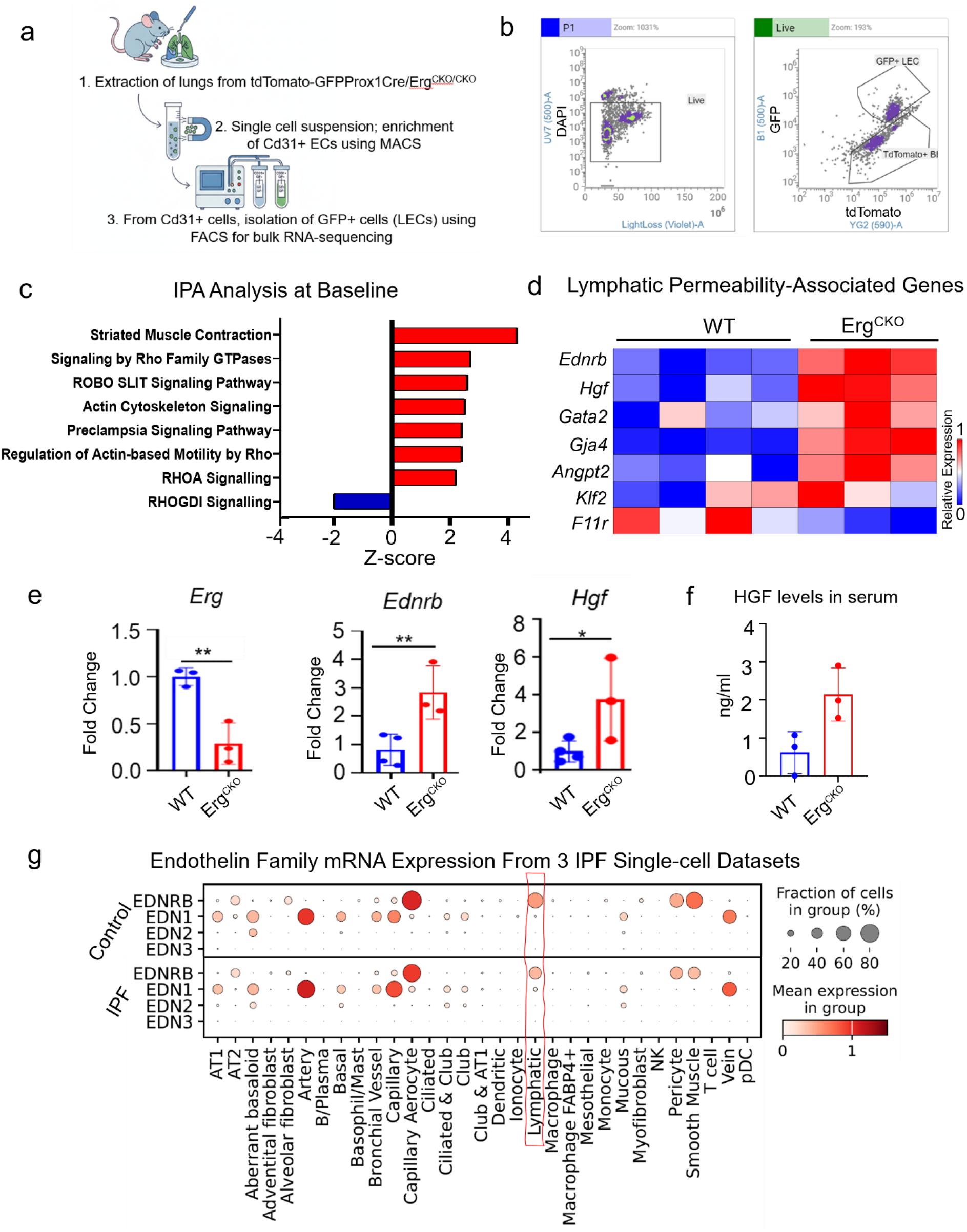
Lymphatic ERG deficiency presents a contractile gene signature and upregulates *Ednrb* and *Hgf*. **a)** Schematic experimental workflow for bulk RNA-sequencing of pulmonary LECs isolated from Erg-CKO-GFP and WT-GFP mice. **b)** Representative FACS gating strategy for sorting live, single GFP+ pulmonary LECs versus tdTomato BECs. **c)** Ingenuity Pathway Analysis (IPA) of bulk RNA-seq data showing significantly enriched and activated signaling pathways (positive Z-scores, red) and inhibited pathways (negative Z-score, blue) in Erg-CKO LECs compared to WT controls at baseline. d) Heatmap of bulk RNA-seq data illustrating the relative expression of lymphatic permeability-associated genes in sorted pulmonary LECs from WT and Erg-CKO mice (n=3–4). Relative expression scale: low (blue) to high (red). **e)** Quantitative RT-qPCR validation of *Erg*, *Ednrb*, and *Hgf* transcript levels (fold change relative to WT) in sorted pulmonary LECs from WT and Erg-CKO mice (n=3). **f)** ELISA quantification of systemic circulating HGF protein levels (ng/mL) in serum from WT and Erg-CKO mice (n=3). **g)** Dot plot showing single-cell RNA sequencing meta-analysis of endothelin system components (*EDNRB, EDN1, EDN2, EDN3*) across major pulmonary cell types in HCs and IPF lungs from three integrated human datasets, highlighting prominent EDNRB expression in lymphatic endothelial cells (red box) and capillary aerocytes. Dot size represents the fraction of expressing cells (%), and color intensity indicates mean expression levels.

### Pharmacological inhibition of EDNRB reverses enhanced lymphatic drainage and protective phenotype in Erg-CKO mice

To directly test whether the upregulation of EDNRB drives the functional advantages seen in Erg-CKO mice, we pharmacologically inhibited EDNRB using its selective antagonist BQ-788. We first evaluated baseline fluid clearance using FITC-dextran drainage assay (**Fig. 6a**). Mice received localized *i.t.* administration of BQ-788 (500 pmol/mouse) or vehicle prior to FITC-dextran instillation, followed by *ex vivo* imaging of the draining mLNs (**Fig. 6a**). As expected, vehicle-treated Erg-CKO mice displayed significantly higher lymphatic drainage compared to WT controls, as evidenced by elevated fluorescence intensity in mLNs (***p<0.001; **Fig. 6b**). However, treatment with BQ-788 eliminated this advantage, reducing tracer transport in Erg-CKO mice down to baseline WT levels (***p<0.001; **Fig. 6b**). We next investigated whether blocking EDNRB altered early acute lung injury dynamics following bleomycin administration (**Fig. 6c**). BALF was collected at 7dpi to evaluate cellular influx and barrier leakage (**Fig. 6c**). While vehicle-treated Erg-CKO mice exhibited a marked reduction in total number of inflammatory cells compared to WT controls, daily intraperitoneal (*i.p.*) administration of BQ-788 fully reversed this protection (****p<0.0001; **Fig. 6d**). Inflammatory cell counts in BQ-788-treated Erg-CKO mice returned to levels comparable to injured WT mice (**Fig. 6d**). Similarly, total BALF protein concentration was significantly suppressed in vehicle-treated Erg-CKO mice (**p<0.01 vs WT) but increased significantly upon EDNRB inhibition (*p<0.05; **Fig. 6d**). Finally, we assessed whether EDNRB blockade would compromise anti-fibrotic protection in Erg-CKO mice at 14dpi. Histological evaluation via Masson’s Trichrome staining revealed that vehicle-treated Erg-CKO mice maintained preserved parenchymal architecture and minimal scarring following bleomycin injury (**Fig. 6e**). In contrast, treatment with BQ-788 abrogated this protective phenotype, leading to dense ECM deposition, severe alveolar thickening, and widespread fibrotic remodeling in Erg-CKO lungs (**Fig. 6e**). Quantitative morphometry confirmed a corresponding increase in the tissue-to-airspace ratio in BQ-788-treated Erg-CKO mice, demonstrating a reversal toward an injured, WT-like phenotype (****p<0.0001; **Fig. 6f**). Notably, fibrotic severity in BQ-788-treated WT mice was comparable to vehicle-treated WT controls at 14dpi (**Fig. 6f**); however, EDNRB antagonism resulted in substantial mortality, with 50% of the WT mice succumbing to severe lung injury between 10 and 13dpi (Data not shown). Taken together, these functional, biochemical, and histological experiments establish EDNRB as a downstream mediator through which lymphatic ERG deficiency accelerates fluid clearance, attenuates early acute inflammation, and protects against the progression of pulmonary fibrosis.

**Figure 6.**
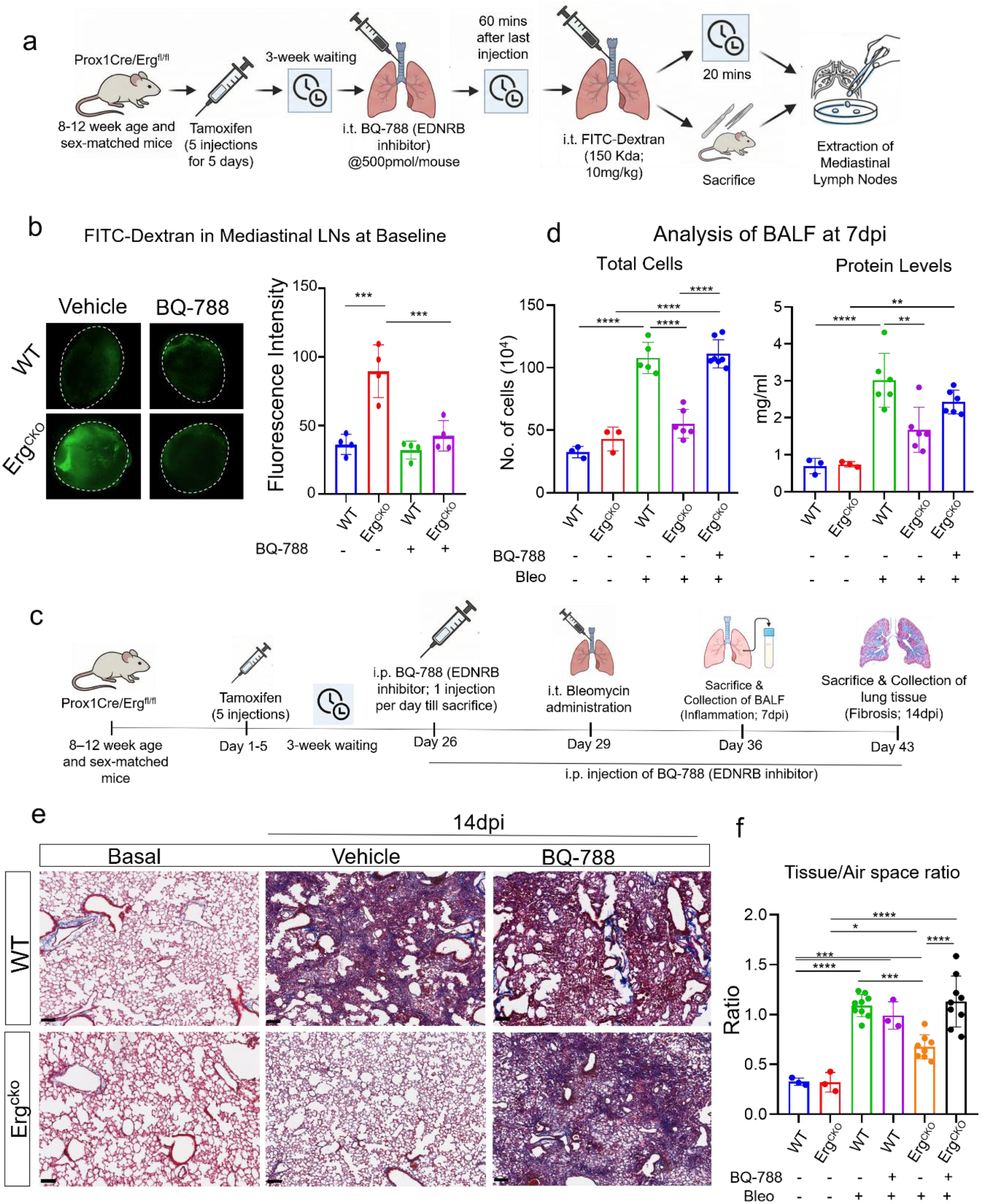
EDNRB signaling is required for enhanced lymphatic drainage and protective phenotype in Erg-CKO mice. **a)** Schematic experimental outline for *i.t.* administration of the selective EDNRB antagonist BQ-788 (500 pmol/mouse) followed 60 minutes later by *i.t.* instillation of FITC-Dextran (150 kDa; 10 mg/kg) to evaluate lymphatic drainage transport to mLNs (harvested 20 minutes post-dextran instillation). **b)** Representative fluorescence images (left; dashed outlines) and quantitative fluorescence intensity analysis (right) of FITC-Dextran accumulation in draining mLNs of vehicle- or BQ-788-treated wild-type (WT) and Erg-CKO mice under baseline conditions (n=4). **c)** Schematic experimental timeline for *i.p.* BQ-788 administration (daily injections starting 3 days prior to bleomycin instillation) and subsequent assessment of acute inflammation at 7dpi (BALF collection) and fibrotic remodeling at 14dpi (lung histology). **d)** Analysis of BALF at 7dpi showing total inflammatory cell counts (10^4^ cells; left) and total protein concentrations (mg/mL; right) across vehicle-treated WT, vehicle-treated Erg-CKO, and BQ-788-treated Erg-CKO mice subjected to bleomycin injury (n=5–7). **e)** Representative images of Masson’s trichrome staining of lung tissue sections from WT and Erg-CKO mice under baseline conditions and at 14dpi following daily treatment with vehicle or BQ-788, illustrating the complete reversal of the protective phenotype in BQ-788-treated Erg-CKO lungs. Scale bars = 100µm. **f)** Morphometric quantification of parenchymal remodeling expressed as the tissue-to-airspace ratio across experimental groups at baseline and 14dpi (n = 3–9).

### ERG deficiency drives a zipper-to-button junctional transition in LECs through EDNRB activation

To determine the structural mechanism behind the enhanced fluid transport in Erg-CKO mice, we examined the intercellular junctional architecture of pulmonary lymphatic vessels (*in vivo*) and cultured human pulmonary LECs (*in vitro*). We first visualized VE-cadherin distribution on 300µm precision cut lung sections (PCLSs) from WT and Erg-CKO GFP-reporter mice. High-resolution confocal imaging revealed distinct junctional patterns between the two genotypes. WT collector lymphatics displayed continuous, linear VE-cadherin staining along cell-cell borders (green arrowheads), characteristic of mature, continuous zipper-like junctions (**Fig. 7a**); whereas, Erg-CKO collector lymphatics showed a shift toward discontinuous, oakleaf-like VE-cadherin patterns with interjunctional gaps (red arrowheads), representative of button-like junctions (**Fig. 7a**). These structural changes reveal that collecting lymphatics in Erg-CKO mice acquire discontinuous, button-like junctions, consistent with the augmented drainage observed *in vivo*. To confirm whether this junctional transition is cell-intrinsic and mediated by EDNRB, we performed siRNA-mediated knockdowns in primary human LECs. Cells were treated with siSCR, siERG, or siERG in combination with the EDNRB antagonist BQ-788, followed by IF staining for VE-cadherin. The control siSCR-treated cells showed intact, continuous monolayers with VE-cadherin localization around the entire cell perimeter, whereas siERG treatment triggered noticeable junctional remodeling characterized by serrated borders, localized VE-cadherin retraction, and clear intercellular gap formation (red arrowheads; **Fig. 7b**). Further, pharmacological inhibition of EDNRB blunted this gap formation noticeably and partially restoring the more continuous junctional pattern along cell boundaries, unlike siERG only treated cells (**Fig. 7b**).

**Figure 7.**
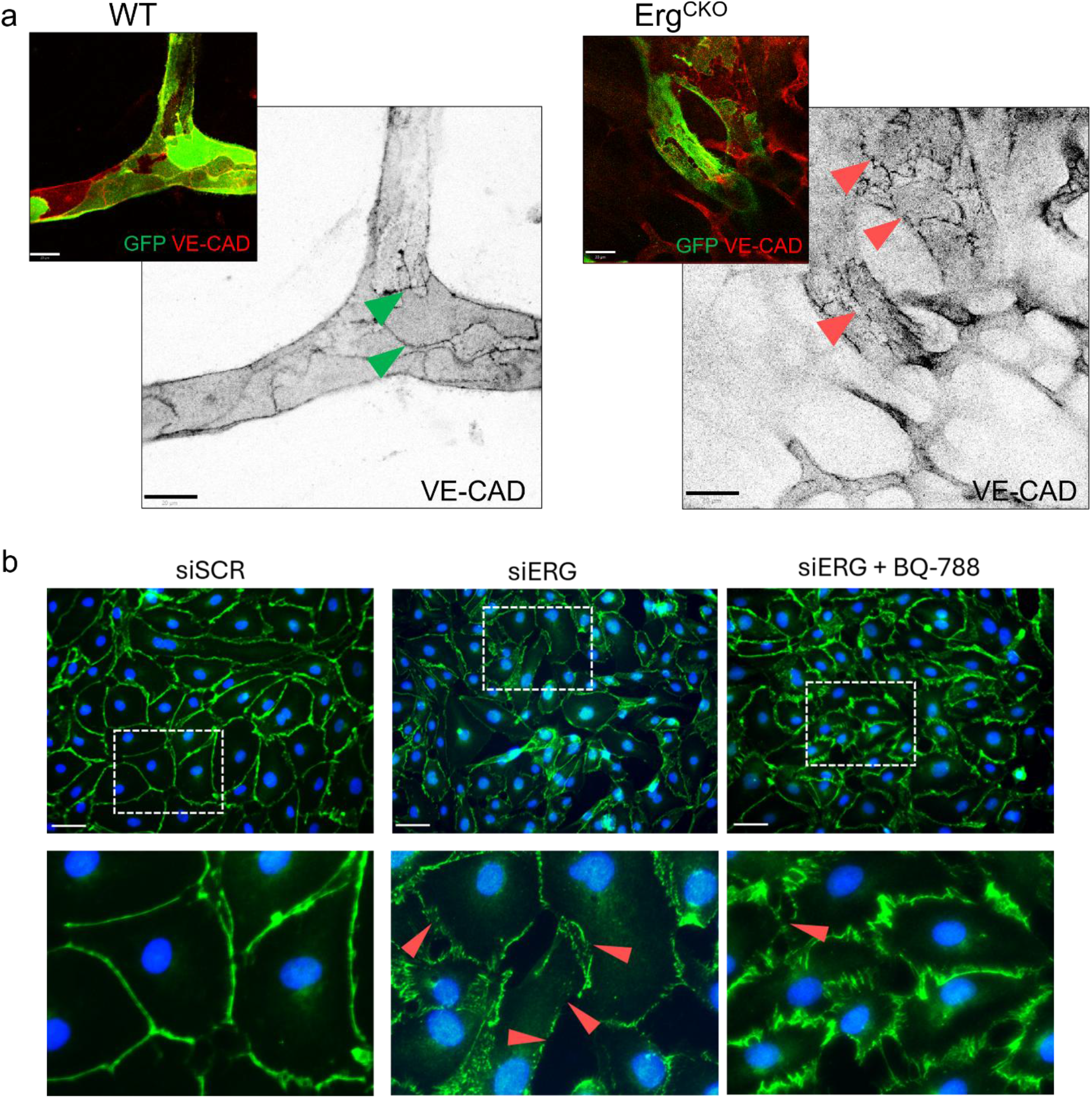
ERG deficiency drives a zippered-to-buttoned junctional transition in LECs via EDNRB activation. **a)** High-resolution confocal microscopy images of pulmonary lymphatic vessels in lung tissue sections from WT (left) and Erg-CKO mice (right) stained for endogenous lymphatic GFP (green) and VE-Cadherin (red), alongside inverted grayscale panels of the VE-Cadherin channel. Note the continuous, linear zipper-like cell–cell junctions in WT lymphatics (green arrowheads) versus the discontinuous, punctate button-like absorptive junctional patterns in Erg-CKO lymphatics (red arrowheads). Scale bars = 20µm. **b)** Representative IF confocal micrographs of cultured human pulmonary LECs transfected with siSCR (left) or siERG in the absence (middle) or presence (right) of the selective EDNRB antagonist BQ-788, stained for VE-Cadherin (green) and nuclei (DAPI, blue). Lower panels show high-magnification views of the boxed regions, highlighting the formation of inter-endothelial gaps and discontinuous button-like junctional remodeling following ERG knockdown (red arrowheads), which is effectively rescued/reversed upon pharmacological EDNRB inhibition with BQ-788. Scale bars = 20µm.

Together, these results demonstrate that ERG deficiency is associated with a structural transformation from continuous zippered junctions to absorptive buttoned junctions in LECs *in vivo*. Furthermore, the reversal of this phenotype by BQ-788 *in vitro* suggests that EDNRB activation may drive this junctional remodeling, providing a plausible structural mechanism for the enhanced drainage and anti-fibrotic protection seen in Erg-CKO lungs.

## DISCUSSION

This study reveals a novel regulatory axis in pulmonary lymphatic biology, establishing the transcription factor ERG as a critical gatekeeper of adult lymphatic quiescence. Loss of ERG upregulates EDNRB to enhance lymphatic clearance and attenuate bleomycin-induced pulmonary fibrosis. These findings position the pulmonary lymphatic network as an active, targetable niche in fibroproliferative lung disease and highlight the cell-type-and organ-specific functions of ERG. In the blood vascular endothelium, ERG acts as a master regulator of structural integrity. Constitutive ERG expression preserves endothelial quiescence by promoting VE-cadherin transcription and repressing inflammatory cascades, whereas its loss leads to vascular leakage, inflammation, and pathological remodeling (7, 8, 10).

Although LECs and BECs share a common endothelial lineage, they serve fundamentally distinct physiological purposes (16). Interestingly, the enhanced lymphatic drainage we observed appears to be a direct consequence of increased vessel permeability, mechanistically mirroring the hyperpermeability seen in blood vessels following endothelial ERG loss. This suggests how similar cellular perturbation can produce distinct physiological outcomes depending on the vascular bed: while increased permeability in the blood vasculature promotes plasma leakage and fuels inflammation, heightened permeability in lymphatics accelerates the uptake and clearance of interstitial fluid and inflammatory exudates, thereby resolving tissue edema. This concept is well-supported by studies demonstrating that augmenting lymphatic drainage facilitates the resolution of acute inflammation and prevents tissue remodeling (17–20). Whether this protective phenotype represents an active adaptive mechanism remains an important question. Because lymphatic ERG expression naturally declines in both injured mouse lungs and human IPF tissue, ERG downregulation may serve as an evolutionarily conserved trigger to enhance fluid clearance during parenchymal injury.

Further investigation led us to the G-protein–coupled receptor EDNRB as the central downstream mediator coordinating these functional and structural adaptations. In the pulmonary vascular bed, EDNRB is predominantly expressed on endothelial subsets, including lymphatic endothelial cells and specialized capillary aerocytes (Cap2), where it regulates endothelial nitric oxide release, endothelin-1 clearance, and localized cytoskeletal dynamics (21, 22). In our proposed model, upregulation of EDNRB following ERG loss may facilitate the structural transition of VE-cadherin cell-cell contacts from continuous zippers into absorptive button-like formations that accelerate interstitial clearance. Such a junctional shift is a well-established mechanism that lowers entry resistance to rapidly clear protein-rich edema (5, 19, 23). While further *in vivo* structural studies will be needed to definitively confirm this remodeling, our finding that BQ-788 reverses intercellular gap formation in siERG-treated human pulmonary LECs *in vitro* supports the possibility that EDNRB signaling contributes to this junctional plasticity.

This capacity of LECs to actively tune their transport machinery aligns with emerging literature on mechanical and receptor-mediated lymphatic control. For instance, activating the mechanosensitive ion channel Piezo1 via Yoda-1 enhances meningeal lymphatic drainage to alleviate excessive cerebrospinal fluid accumulation (24). Similarly, activation of PAR-1 in LECs leads to junctional remodeling to enhance lymphatic drainage during acute lung injury (19). Our transcriptomic findings support this paradigm. Pathway analysis of Erg-CKO LECs revealed positive activation of cascades governing actomyosin contractility, Rho family GTPases, and actin cytoskeleton dynamics. Together, these findings link ERG deficiency and downstream EDNRB upregulation to an integrated program of cytoskeletal contraction and junctional reorganization, optimizing pulmonary lymphatics for rapid fluid uptake. Furthermore, pharmacological blockade of EDNRB with BQ-788 abolished the enhanced drainage in Erg-CKO mice and exacerbated fibrotic damage. Moreover, BQ-788 worsened fibrosis and increased mortality in WT mice, underscoring the crucial role of EDNRB signaling in protecting against lung fibrosis. In addition to facilitating lymphatic drainage, EDNRB expression characterizes cap2 cells and is involved in ET-1 clearance (21, 22). Inhibition of EDNRB signaling destabilizes the alveolar–capillary barrier, exacerbates gas-exchange impairment, and increases mortality.

From a translational perspective, EDNRB represents a viable pharmacological target to combat edematous and fibroproliferative lung diseases. While classical pulmonary fibrosis therapies focus primarily on fibroblast inhibition, boosting lymphatic clearance offers a disease-modifying strategy to decompress the injured interstitium and prevent aberrant tissue remodeling. Selective EDNRB agonism via Sovateltide (IRL-1620), which is currently in Phase III clinical trials with an established safety profile, has shown potent tissue-protective, anti-apoptotic, and hemodynamic benefits across ischemic and vascular injury models (25, 26). Beyond direct mechanical clearance, elevated HGF expression in *Erg*-deficient LECs provides a vital paracrine defense. Given HGF’s known anti-fibrotic activity, we hypothesize that secreted HGF binds c-Met on adjacent alveolar epithelial cells to counter TGF-β1–driven fibrotic transformation and promote epithelial repair (27–29), acting synergistically with EDNRB-mediated drainage to reinforce the anti-fibrotic phenotype, but future investigations will be required to address this.

While our findings demonstrate the therapeutic potential of targeting pulmonary lymphatics, several translational and mechanistic questions remain. First, because ERG is a master transcription factor with broad biological roles, its deletion triggers complex downstream effects beyond EDNRB induction, including the downregulation of canonical lymphatic identity genes alongside the activation of contractile programs and HGF secretion. Consequently, direct systemic manipulation of ERG carries broad pleiotropic risks, underscoring why targeting specific downstream effectors like EDNRB provides a safer and more tractable therapeutic approach. Second, while our findings clearly show that EDNRB regulates lymphatic drainage, whether it is the sole driver of the Erg-CKO protective phenotype remains to be defined. While our drainage assay directly reflects a lymphatic-specific function, overall fibrotic protection involves multiple cell types. Because pharmacological BQ-788 treatment targets not only lymphatics but also other EDNRB-expressing populations, including capillary aerocytes and vascular smooth muscle cells, we cannot exclude the possibility that non-lymphatic vascular stress contributed to the worsened pathology. Hence, future studies utilizing lymphatic-specific Ednrb-CKO and Erg/Ednrb double-knockout genetic mice lines will be necessary to completely isolate these cell-autonomous mechanisms *in vivo*. Third, while our study established pathway necessity through selective BQ-788 antagonism in Erg-CKO mice, clinical translation will require direct pharmacological stimulation. Evaluating selective EDNRB agonists, such as Sovateltide (IRL-1620), across multiple therapeutic windows represents a vital next step for preclinical testing.

In conclusion, our study identifies the ERG-EDNRB axis as a vital regulator of pulmonary lymphatic function and tissue resilience. We demonstrate that the loss of ERG primes the lymphatic system for superior fluid and macromolecule clearance, effectively mitigating the devastating effects of fibrotic lung injury. These findings underscore the therapeutic potential of targeting EDNRB signaling, a pathway with existing clinical precedents, to enhance lymphatic drainage as a regenerative strategy in the fight against pulmonary fibrosis and other forms of respiratory diseases.

## METHODS

### Sex as a biological variable

Our study examined both male and female human lung tissue specimens and cell donors. For animal experiments, age-matched male and female adult mice (8–10 weeks of age) were allocated to experimental and control cohorts. Although preliminary responses to bleomycin and baseline lymphatic phenotypes appeared comparable between sexes, individual cohorts were not statistical powered to assess sex as a biological variable or detect subtle sex-specific interaction effects. Consequently, data from male and female animals were pooled, and potential sex-dependent differences remain to be formally investigated.

### Human Tissue Specimens and Primary Cell Isolation

Human lung tissues from 4 HCs and 6 patients with IPF were obtained from the University of Michigan. For primary human LEC isolation, lungs were minced and enzymatically digested in endothelial cell basal medium MV2 (Cat. C-22022; PromoCell) without supplements containing Collagenase II (0.1% w/v; Cat. 17101015, Gibco), Collagenase IV (0.25% w/v; Cat. NC9919937, Worthington Biochemical Corporation), and DNase 1 (1µl/ml; Cat. 4536282001, Roche) for 1 hour at 37°C with constant rotation. The digest was passed through a 100*μ*m cell strainer twice followed by passing through 40*μ*m cell strainer twice. Cells were grown in Endothelial Cell Growth Medium MV2 supplemented with growth factors (Cat. #C-22121, PromoCell) for few days. Later, endothelial cells were enriched using anti-CD31 magnetic dynabeads (Cat. 11-155-D, Invitrogen). Following bead detachment, pure LECs were isolated by positive selection using an anti-human PDPN-PE antibody (Clone REA446, Cat. 130-117-687, Miltenyi Biotec) coupled with anti-PE magnetic MicroBeads (Cat. 130-048-801, Miltenyi Biotec) and propagated in complete MV2 medium.

### Mice and Tamoxifen-Induced Cre Recombination

Inducible lymphatic-specific *Erg* knockout reporter mice (Erg-CKO) were generated by crossing *Erg*^fl/fl^ mice (possessing *loxP* sites flanking exon 6) with Prox1CreERT2 and Rosa26-mTmG reporter lines (*Erg*^fl/fl^*Prox1*-CreERT2+tdTomato+) as described earlier (14). *Erg*^+/+^*Prox1*-CreERT2^+^tdTomato^+^ littermates served as WT controls. Gene excision was induced in 8-to 10-week-old adult mice by *i.p.* injection of tamoxifen (75 mg/kg in corn oil; Cat. T5648, Sigma-Aldrich) once daily for 5 consecutive days. Experiments were initiated 21 days after the final injection to ensure complete Cre-mediated recombination and drug clearance.

### Bleomycin Model of Pulmonary Fibrosis

Pulmonary injury and fibrosis were induced by a single *i.t.* instillation of bleomycin sulfate (0.05 U per mouse in 50*μ*L sterile PBS; Cat. 1076308, Sigma) under isoflurane anesthesia. Control mice received an equivalent volume of sterile PBS (mentioned as baseline in the text). Lungs, BALF, and blood were collected at defined acute inflammatory (3 and 7dpi) and fibrotic (14dpi) endpoints.

### *In Vivo* Pharmacological EDNRB Inhibition

To block EDNRB *in vivo*, the selective antagonist BQ-788 (Cat. HY-15894A, MedChemExpress) was dissolved in sterile physiological saline. For acute drainage assays, BQ-788 (500pmol/mouse) or vehicle saline was instilled *i.t.* 30 minutes prior to fluorescent tracer delivery. For chronic bleomycin studies, mice received daily *i.p.* injections of BQ-788 (1 mg/kg/day) starting 4 days prior to bleomycin instillation and continuing daily until harvest at 7dpi or 14dpi. Control cohorts received equal volumes of vehicle saline.

### BALF Analysis and Pulmonary Edema

BALF was collected from mouse lungs following standard protocols (30). Briefly, tracheas were exposed and cannulated with a 20-gauge catheter, and lungs were lavaged three times with 1mL ice-cold PBS. The obtained BALF was centrifuged at 1500rpm for 15 minutes at 4°C. Supernatants were collected to measure total protein extravasation via BCA Protein Assay (Cat. 23227, Thermo Fisher Scientific) and Interleukin-6 (IL-6) levels via mouse IL-6 ELISA (Cat. ARG83349, Arigo Biolaboratories Corp.). Cell pellets were resuspended in PBS, and total inflammatory cell counts were determined using a hemocytometer. To quantify pulmonary edema, superior right lung lobes from unlavaged mice were weighed immediately upon harvest (wet weight), dried at 65°C for 48 hours, and reweighed (dry weight) to calculate the wet-to-dry weight ratio.

### Histopathology, Whole-Slide Morphometry, and Hydroxyproline Assay

Lungs were inflation-fixed with 4% paraformaldehyde (PFA) for 24 hours, embedded in paraffin, and cut into 5µm sections. Collagen deposition and architectural distortion were visualized with Masson’s Trichrome staining. Whole-slide digital scans were acquired using a PhenoImager HT System slide scanner from Akoya Biosciences, and parenchymal consolidation (tissue-to-airspace ratio) was quantified across the entire lung area using QuPath software. Total lung collagen content was determined biochemically from left lung homogenates using a commercial Hydroxyproline Assay Kit (Cat. MAK569, Sigma-Aldrich) according to the manufacturer’s protocol and expressed as *μ*g of hydroxyproline/10mg of lung tissue.

### *In Vivo* Pulmonary Lymphatic Drainage Assay

Macromolecular lymphatic clearance was quantified by *i.t.* instillation of high-molecular-weight FITC-dextran (150 kDa; 10mg/kg in sterile PBS, 50*μ*L total injection volume; Cat. FD150S, Sigma-Aldrich) into anesthetized WT and Erg-CKO mice at baseline or 7dpi. After 20 minutes of *in vivo* transport, draining mLNs were dissected and immediately imaged *ex vivo* on the green fluorescence channel using an Olympus CKX53 inverted microscope equipped with a fluorescence module and a Lumenera Infinity 3 digital camera. Total fluorescence integrated density in mLNs was quantified using ImageJ. Background autofluorescence was subtracted using non-instilled control mice.

### PCLS and IF Staining

For junctional analysis, mouse lungs were inflated with 2% low-melting-point agarose in PBS, cooled on ice, and sliced into 300µm thick sections using a vibratome by Precisionary Instruments and were stained for junctions as defined earlier (23). Paraffin sections (5µm) from human and mouse lungs were deparaffinized, subjected to heat-induced antigen retrieval in citrate buffer (pH 6.0), permeabilized with 0.2% Triton X-100, and blocked in 5% donkey serum. Tissues and PCLS were incubated overnight at 4°C with primary antibodies: anti-PDPN (1:200; Clone NZ-1.3, Cat. 14-9381-82, Invitrogen), anti-ERG (1:200; Clone A7L1G, Cat. 97249, Cell Signaling Technology), anti-Col1a1 (1:200; Clone E8F4L, Cat. 72026, Cell Signaling Technology), anti-GFP (1:500; Cat. NC0407892, Aves Labs), and anti-VE-Cadherin (1:200; Cat. AF1002, R&D Systems). Samples were washed and incubated with Alexa Fluor-conjugated secondary antibodies (1:1000; Thermo Fisher Scientific) and DAPI (1µg/mL). PCLSs images were acquired using a confocal microscope, Zeiss LSM 700, whereas paraffin sections (both human and mouse) were imaged using Akoya Biosciences PhenoImager HT System.

### Quantitative Image Analysis of Human Lung Tissue

Quantitative image analysis was performed using QuPath (v0.5.1), where the whole sections were utilized as region of interest (ROI) to keep the analysis unbiased. Total cellularity was determined via DAPI-based nuclear detection using the Cell Detection command with a requested pixel size of 0.5µm, a minimum nuclear area of 15 µm², and a maximum area of 200 µm². ERG-positive (ERG^+^) cells were identified using a Single Measurement Classifier where cells were classified as ERG+ if the “Nucleus: ERG Mean intensity” exceeded a threshold of 350, determined via bimodal histogram distribution analysis. LECs were identified as cells with PDPN+ object classification defined by the overlap of PDPN staining within the defined cell boundary. A spatial hierarchy was subsequently resolved (Analyze > Objects > Resolve hierarchy) to nest cell detections within their respective parent annotations, allowing for the binary categorization of ERG^+^ and ERGˉ cells located within or outside LECs (annotated by PDPN+ LECs). Finally, the fraction of ERG+ cells within LECs against total number of LECs were quantified. The representation of the object classifiers is shown in **Supplementary Figure 1**.

### *In Vitro* siRNA Transfection and Cell Culture Assays

For transfection experiments, isolated primary human pulmonary LECs (passages 3–6) were transfected with 25nM ON-TARGETplus non-targeting control siRNAs (siSCR; Cat. D-001810-01-05, Horizon) or *ERG*-targeting siRNA (siERG; Cat. J-003886-09-0002, Horizon) using lipofectamine RNAiMAX transfection reagent (Cat. 13778075, Thermo Fisher Scientific). Target gene knockdown efficiency was verified by RT-qPCR (**Supplementary Fig. 4a**). The knockdown in rescue experiments, cells were co-treated with BQ-788 (10µM) or vehicle for 12 hours. Confluent monolayers were fixed with 4% PFA, permeabilized with 0.1% Triton X-100, blocked in 5% donkey serum, and stained for VE-Cadherin (1:200; Cat. AF1002, R&D Systems) and DAPI.

### Primary Mouse LEC Isolation and FACS Sorting

Lungs from WT and Erg-CKO reporter mice were digested in Collagenase II (0.1% w/v) and IV (0.25% w/v) for 45 minutes at 37°C. Single-cell suspensions were incubated with anti-CD31 magnetic MicroBeads (Cat. 130-097-418, Miltenyi Biotec) to enrich endothelial cells via autoMACS columns. Enriched cells were resuspended in sorting buffer containing DAPI and sorted on a BD FACSAria II SORP cell sorter (BD Biosciences) using a 100µm nozzle. Gating on forward/side scatter and DAPI exclusion isolated live single cells, followed by sorting of pure GFP+ LECs away from tdTomato+ BECs directly into lysis buffer.

### Bulk RNA-Sequencing and Transcriptomic Analysis

Total RNA was extracted from sorted mouse LECs (WT vs. Erg-CKO at baseline, 7dpi, and 14dpi) and cultured human LECs (siSCR vs. siERG) using the RNeasy Plus Micro Kit (Cat. 74034, Qiagen). Due to low cell yields (∼500–1,000 sorted LECs per mouse sample), RNA samples approached capillary electrophoresis detection thresholds (Agilent 2100 Bioanalyzer); integrity was confirmed by evaluating electropherograms for distinct 18S and 28S peaks (RIN > 5.0). Low-input cDNA libraries were prepared and sequenced on an Illumina NextSeq/NovaSeq platform (paired-end). Reads were aligned to the mm10 mouse or GRCh38 human reference genomes using STAR (v2.7.9a). Gene-level counts were quantified using featureCounts (v1.6.2). Normalization, sample clustering via PCA, and differential expression were conducted in R (v4.1.2) using DESeq2 (v1.23.10), defining significance at Benjamini-Hochberg FDR q<0.05 and fold change ≥ 1.5. Functional pathway enrichment and activation Z-scores were computed using Ingenuity Pathway Analysis (IPA; Qiagen).

### Quantitative Real-Time PCR (RT-qPCR) and Serum ELISA

cDNA was synthesized using the High-Capacity cDNA Reverse Transcription Kit (Cat. 4374966, Fisher Scientific). Quantitative PCR was performed using PowerUp SYBR Green Master Mix (Cat. A25742, Thermo Fisher Scientific) on a Quantstudio 3 thermocycler (appliedbiosystems). Target gene expression was normalized to *Gapdh* using the 2−ΔΔCt method. Primer sequences are listed in **Supplementary Table 1**. For circulating systemic HGF measurements, whole blood was collected by cardiac puncture, allowed to clot at room temperature for 30 minutes, and centrifuged at 2,000×*g* for 10 minutes at 4°C to collect serum. Circulating HGF was quantified using the Mouse HGF ELISA Kit (Cat. EMHGF, Thermo Fisher Scientific).

### Human Single-Cell RNA-Seq Meta-Analysis

Public single-cell RNA-sequencing datasets from healthy human donor lungs and IPF patients (Habermann, Morse, and Adams cohorts) were integrated. Major lung cell types, including lymphatic endothelial cells, capillary aerocytes, general capillaries, arteries, veins, and stromal subsets, were clustered and annotated. Transcript abundance and cell-type distribution for *ERG* and endothelin system members (*EDNRB, EDN1, EDN2, EDN3*) were visualized using dot plots displaying expression percentage and mean intensity.

### Statistical Analysis

Data are expressed as mean±SD. Comparisons between two groups were analyzed using two-tailed unpaired Student’s *t*-test. Comparisons among three or more groups were evaluated using one-way or two-way ANOVA followed by Tukey’s post hoc multiple comparisons test. Statistical analyses were performed in GraphPad Prism (v9.0/10.0), and significance was set at p<0.05.

### Study approval

All animal experimental procedures were reviewed and approved by the Boston University Institutional Animal Care and Use Committee (IACUC Protocol 201800144_TR01) and conformed to the NIH *Guide for the Care and Use of Laboratory Animals*. Human lung tissue procurement and experimental protocols were approved by the University of Michigan Institutional Review Board (IRB) for Human Studies in compliance with all relevant ethical guidelines under IRB HUM00105694. Written informed consent was obtained from all patients or their legal surrogates prior to tissue collection.

## Data availability

All raw and processed bulk RNA-sequencing data generated in this study have been deposited in the NCBI Gene Expression Omnibus (GEO) database. These data are available under accession numbers GSE344150 (human LECs) and GSE345222 (mouse LECs). Public human single-cell RNA-sequencing datasets analyzed in this study are available in GEO under accessions GSE135893 (Habermann et al.), GSE128033 (Morse et al.), and GSE136831 (Adams et al.). All underlying source data for graphs and figures are provided in the Supporting Data Values file.

## Author contributions

**A.N.**: Conceptualization, methodology, validation, formal analysis, investigation, data curation, writing–original draft, visualization, project administration. **T.M.**: Methodology, investigation, data curation, validation. **X.Q.**: Formal analysis (specifically single-cell RNA-seq meta-analysis), software, visualization. **A.C.**: Methodology, investigation. **A.A.R.**: Methodology, investigation. **G.L.**: Conceptualization, resources, human specimen acquisition. **M.T.**: Conceptualization, supervision, funding acquisition, writing– review & editing, resources. All authors participated in manuscript preparation and provided final approval of the submitted work.

## Acknowledgements

We thank Dr. Anna C. Belkina and Shari Brezinsky at the Boston University Flow Cytometry Core Facility for their technical expertise and assistance with cell sorting. We also acknowledge Dr. Yuriy Alekseyev and Adam Gower at the Boston University Microarray and Sequencing Resource Core Facility for their expert guidance in RNA-sequencing experimental design, library preparation, and bioinformatic analysis.

## References

1. Martinez FJ, Collard HR, Pardo A, Raghu G, Richeldi L, Selman M, et al. Idiopathic pulmonary fibrosis. Nature Reviews Disease Primers. 2017;3(1):17074.

2. Richeldi L, Collard HR, and Jones MG. Idiopathic pulmonary fibrosis. The Lancet. 2017;389(10082):1941–52.

3. Matthay MA, Zemans RL, Zimmerman GA, Arabi YM, Beitler JR, Mercat A, et al. Acute respiratory distress syndrome. Nature Reviews Disease Primers. 2019;5(1):18.

4. Alitalo K. The lymphatic vasculature in disease. Nature Medicine. 2011;17(11):1371–80.

5. Baluk P, Fuxe J, Hashizume H, Romano T, Lashnits E, Butz S, et al. Functionally specialized junctions between endothelial cells of lymphatic vessels. The Journal of experimental medicine. 2007;204(10):2349–62.

6. Trivedi A, and Reed HO. The lymphatic vasculature in lung function and respiratory disease. Frontiers in Medicine. 2023;10:1118583.

7. Birdsey GM, Shah AV, Dufton N, Reynolds LE, Osuna Almagro L, Yang Y, et al. The endothelial transcription factor ERG promotes vascular stability and growth through Wnt/β-catenin signaling. Developmental cell. 2015;32(1):82–96.

8. Dryden NH, Sperone A, Martin-Almedina S, Hannah RL, Birdsey GM, Khan ST, et al. The transcription factor Erg controls endothelial cell quiescence by repressing activity of nuclear factor (NF)-κB p65. The Journal of biological chemistry. 2012;287(15):12331–42.

9. Shah AV, Birdsey GM, and Randi AM. Regulation of endothelial homeostasis, vascular development and angiogenesis by the transcription factor ERG. Vascular pharmacology. 2016;86:3–13.

10. Caporarello N, Lee J, Pham TX, Jones DL, Guan J, Link PA, et al. Dysfunctional ERG signaling drives pulmonary vascular aging and persistent fibrosis. Nature communications. 2022;13(1):4170.

11. Adams TS, Schupp JC, Poli S, Ayaub EA, Neumark N, Ahangari F, et al. Single-cell RNA-seq reveals ectopic and aberrant lung-resident cell populations in idiopathic pulmonary fibrosis. Science advances. 2020;6(28):eaba1983.

12. Habermann AC, Gutierrez AJ, Bui LT, Yahn SL, Winters NI, Calvi CL, et al. Single-cell RNA sequencing reveals profibrotic roles of distinct epithelial and mesenchymal lineages in pulmonary fibrosis. Science advances. 2020;6(28):eaba1972.

13. Morse C, Tabib T, Sembrat J, Buschur KL, Bittar HT, Valenzi E, et al. Proliferating SPP1/MERTK-expressing macrophages in idiopathic pulmonary fibrosis. The European respiratory journal. 2019;54(2).

14. Yamashita T, Kaplan U, Chakraborty A, Marden G, Gritli S, Roh D, et al. ERG Regulates Lymphatic Vessel Specification Genes and Its Deficiency Impairs Wound Healing-Associated Lymphangiogenesis. Arthritis & rheumatology (Hoboken, NJ). 2024;76(11):1645–57.

15. Mazzuca MQ, and Khalil RA. Vascular endothelin receptor type B: structure, function and dysregulation in vascular disease. Biochemical pharmacology. 2012;84(2):147–62.

16. Ng CP, Helm CL, and Swartz MA. Interstitial flow differentially stimulates blood and lymphatic endothelial cell morphogenesis in vitro. Microvascular research. 2004;68(3):258–64.

17. Schwager S, and Detmar M. Inflammation and Lymphatic Function. Frontiers in immunology. 2019;10:308.

18. Zhang PH, Han J, Cao F, Liu YJ, Tian C, Wu CH, et al. PCTR1 improves pulmonary edema fluid clearance through activating the sodium channel and lymphatic drainage in lipopolysaccharide-induced ARDS. Journal of cellular physiology. 2020;235(12):9510–23.

19. Chou C, Paredes CC, Summers B, Palmer-Johnson J, Trivedi A, Bhagwani A, et al. The thrombin receptor PAR1 orchestrates changes in lymphatic endothelial cell junction morphology to augment lymphatic drainage during lung injury. Nature Cardiovascular Research. 2025;4(8):964–75.

20. Crossey E, Carty S, Shao F, Henao-Vasquez J, Ysasi AB, Zeng M, et al. Influenza induces lung lymphangiogenesis independent of YAP/TAZ activity in lymphatic endothelial cells. Scientific reports. 2024;14(1):21324.

21. Dupuis J, Goresky CA, and Fournier A. Pulmonary clearance of circulating endothelin-1 in dogs in vivo: exclusive role of ETB receptors. Journal of applied physiology (Bethesda, Md : 1985). 1996;81(4):1510–5.

22. Gillich A, Zhang F, Farmer CG, Travaglini KJ, Tan SY, Gu M, et al. Capillary cell-type specialization in the alveolus. Nature. 2020;586(7831):785–9.

23. Chou C, and Reed HO. Lung Lymphatics in Edema, Inflammation, and Thrombosis. *Arteriosclerosis*, Thrombosis, and Vascular Biology. 2025;45(12):2143–54.

24. Choi D, Park E, Choi J, Lu R, Yu JS, Kim C, et al. Piezo1 regulates meningeal lymphatic vessel drainage and alleviates excessive CSF accumulation. Nature Neuroscience. 2024;27(5):913–26.

25. Gulati A, Agrawal N, Vibha D, Misra UK, Paul B, Jain D, et al. Safety and Efficacy of Sovateltide (IRL-1620) in a Multicenter Randomized Controlled Clinical Trial in Patients with Acute Cerebral Ischemic Stroke. CNS drugs. 2021;35(1):85–104.

26. Leonard MG, and Gulati A. Endothelin B receptor agonist, IRL-1620, enhances angiogenesis and neurogenesis following cerebral ischemia in rats. Brain Research. 2013;1528:28–41.

27. Mizuno S, Matsumoto K, and Nakamura T. Hepatocyte growth factor suppresses interstitial fibrosis in a mouse model of obstructive nephropathy. Kidney International. 2001;59(4):1304–14.

28. Shukla MN, Rose JL, Ray R, Lathrop KL, Ray A, and Ray P. Hepatocyte growth factor inhibits epithelial to myofibroblast transition in lung cells via Smad7. American journal of respiratory cell and molecular biology. 2009;40(6):643–53.

29. Panganiban RA, and Day RM. Hepatocyte growth factor in lung repair and pulmonary fibrosis. Acta pharmacologica Sinica. 2011;32(1):12–20.

30. Narota A, Singh R, Bansal R, Kumar A, and Naura AS. Isolation & identification of anti-inflammatory constituents of Randia dumetorum lamk. fruit: Potential beneficial effects against acute lung injury. Journal of Ethnopharmacology. 2023;301:115759.

